# Mouse IgG2a antibodies specific for the commensal *Streptococcus mitis* show stronger cross-reactivity with *Streptococcus pneumoniae* than IgG1 antibodies

**DOI:** 10.1101/635235

**Authors:** Sudhanshu Shekhar, Rabia Khan, Ata Ul Razzaq Khan, Fernanda Cristina Petersen

## Abstract

Here we show that mouse IgG2a and IgG1 antibodies specific for the commensal *Streptococcus mitis* cross-react with the pathogen *Streptococcus pneumoniae*, although the cross-reactivity conferred by IgG2a is stronger than IgG1 antibodies. These findings may have implications for designing *S. mitis*-based vaccines against pneumococcal infections.

## Introduction

*Streptococcus pneumoniae*, a gram-positive bacterium that resides in the human upper respiratory tract, causes several diseases worldwide, such as pneumonia, sepsis, meningitis, and otitis media.^1^ Due to differences in polysaccharide capsules, *S. pneumoniae* has been classified into more than 90 serotypes, which reflect different antigenicity and geographical distribution.^2^ Current pneumococcal vaccines are effective against only those *S. pneumoniae* serotypes that are included in vaccines.^2^ This underscores the exploration of alternative prophylactic strategies that provide protection against all serotypes. In recent years, *Streptococcus mitis*, a commensal that shares phylogenetic, antigenic, and ecological characteristics with *S. pneumoniae*, has appeared as an interesting vaccine candidate that holds the potential to confer optimal immunity to pneumococcal infections.^3,4^

Although recent studies have indicated a crucial role for IgG antibodies in defense against pneumococcal infections, it is unclear as to how *S. mitis* stimulates the host’s immune system to generate humoral responses.^5^ Emerging evidence reveals that rabbits immunized with *S. mitis* generate IgG antibodies that are reactive with both *S. mitis* and *S. pneumoniae*.^4^ In line with this, we have recently shown that mouse IgG antibodies specific for *S. mitis* exhibited cross-reactivity toward *S. pneumoniae* strains D39 (serotype 2) and TIGR4 (serotype 4).^3^ Murine IgG antibodies are heterogeneous consisting of four subclasses (IgG1, IgG2a, IgG2b and IgG3). Out of which IgG2a and IgG1 antibodies play an important role in pathogen defense by opsonization/complement fixation and immune effector functions, respectively.^6,7^ Furthermore, nasopharyngeal colonization of mice with *S. pneumoniae* triggers a significant rise in the levels of antigen-specific IgG2a and IgG1 antibodies.^8^ It however remains unknown whether *S. mitis*-specific IgG antibody responses that are cross-reactive to *S. pneumoniae* are biased to an IgG subclass, such as IgG2a and IgG1. Therefore, the goal of this study was to explore whether: 1) *S. mitis* induces the production of antigen-specific IgG1 and IgG2a antibodies; and 2) these antibodies cross-react with *S. pneumoniae* serotypes. To accomplish this, we intranasally immunized mice with *S. mitis* and examined the IgG reactivity to *S. mitis* and *S. pneumoniae* by Western blotting and ELISA. Our findings provide new knowledge on *S. mitis* mediated production of IgG isotypes that show cross-reactivity with *S. pneumoniae*. This may be important for understanding commensal bacteria-host interplay and designing commensal-based vaccines against pneumococcal infections.

## Material and methods

### Bacterial culture and lysis

The bacterial strains used in this study included *S. mitis* CCUG31611 (type strain, equivalent to NCTC12261), *S. pneumoniae* D39 (serotype 2), and *S. pneumoniae*TIGR4 (serotype 4). All strains were cultured, harvested, and stored as described previously.^4^ To lyse the bacterial cells, a Precellys Lysing Kit and Homogenizer (Precellys 24, Bertin Instruments) was used as per the manufacturer’s instructions. After homogenization, the bacterial cell lysate was centrifuged at 1,000 *g* for 5 min at 4°C and the supernatant was collected and stored at −80°C for further use. The total protein in the cell lysate was quantified using BCA Protein Assay Kit (ThermoFischer Scientific).

### Mouse immunization

CD1 mice used in this study were females of 6-8 weeks age. These mice were specific pathogen free (SPF) and housed in a Minimal Disease Unit at the animal facility at Oslo University Hospital, Rikshospitalet, Oslo, Norway. All animal experiments were approved by the Norwegian Food Safety Authority, Oslo, Norway (Project license number FOTS – 8481). The mice were intranasally immunized with 5 × 10^7^ colony forming units (CFU) of *S. mitis* in 20 µl of PBS or 20 µl of PBS (sham) for each mouse at days 0, 14, and 21. The mice were euthanized at day 7 after the last immunization to collect the blood via intracardiac injection. The freshly isolated blood was kept at 4°C for 1 hour and then centrifuged at 1000g for 5 minutes. The supernatant sera were collected and preserved at −80°C refrigerator for further analysis.

### Western blotting

As descried previously,^3^ SDS-PAGE-separated proteins were transferred from gradient gels (4–20%) to nitrocellulose membranes using a constant voltage of 100 V for 2 h using the BioRad Midi PROTEAN Western blotting module according to the manufacturer’s instructions (BioRad, CA, USA). The membranes were blocked with blocking solution [Tris-buffered saline (TBS) (pH 7.4) containing 0.1% Tween 20 (Sigma-Aldrich) and 5% BSA] for 1 h at room temperature, washed three times with TBS-Tween, and incubated overnight with antisera at 4°C. The antisera were diluted 1:1000. Following incubation, the membranes were washed and finally incubated with AP conjugated anti-mouse IgG2a or IgG1 secondary antibodies and later developed with the substrate (4-chloro-1-naphthol).

### Whole cell ELISA

To determine antibody levels in mouse sera samples, a whole cell ELISA was used as described previously.^4^ In brief, each well of a 96-well plate (Maxisorb, Nunc, Thermo Scientific) was coated overnight with 100 µl of bacterial suspension (OD600 0.5), which was washed and then fixed with 10% formalin. The plate was washed and blocked with a blocking buffer (PBS + 0.05% Tween + 1% BSA) and incubation for 1 h at 37°C. Antisera were diluted (1:100) and added to wells in duplicate, incubated for 2 h at room temperature before addition of the anti-mouse IgG2a/HRP or anti-IgG1/HRP secondary antibody (1:10,000) followed by incubation for 2 h at room temperature. The plates were washed and 100 µl of TMB substrate (ThermoFisher Scientific, Rockford, IL, USA) was added to each well. The plate absorbance was measured by reading the plates at 450 nm using a Multi Mode reader (BioTek™ Cytation™ 3; ThermoFischer Scientific).

### Statistics

Unpaired Student’s *t* test was used to compare two groups of mice (GraphPad Prism Software, version 8, Graph Pad, San Diego, CA, USA). A p value less than 0.05 was considered significant.

### Results

Our Western blotting showed that serum IgG2a and IgG1 antibodies from the *S. mitis*-immunized mice cross-reacted with multiple proteins of *S. pneumoniae* D39 and TIGR4. This was reflected by several visible bands that were between 25 to 75 kD (Figure 1A). Two prominent bands were present at around 250 and 25 kD in the membrane lane loaded with *S. mitis*, but not *S. pneumoniae* serotypes (Figure 1A). Compared to IgG2a, IgG1 antibodies from the immunized mice showed weaker reactivity toward *S. mitis* and *S. pneumoniae* (Figure 1A). However, IgG2a and IgG1 antibodies from the mice inoculated with PBS (control mice) gave rise to few faint bands (Figure 1A). Of note, our data also reflected similar findings in the nasal wash and bronchoalveolar lavage of immunized mice (data not shown). We further measured the levels of serum IgG2a and IgG1 antibodies reactive to *S. pneumoniae* using a whole cell ELISA assay. The *S. mitis*-immunized mice displayed significantly higher levels of IgG2 antibodies reactive to *S. mitis* and *S. pneumoniae* D39 and TIGR4 than IgG1 antibodies (Figure 1B). Taken together, these findings reveal that following mucosal immunization with *S. mitis*, mice generate enhanced IgG2a and IgG1 antibody responses that cross-react with *S. pneumoniae* serotypes, although the IgG2a responses are stronger than IgG1 responses.

**Figure 1.**
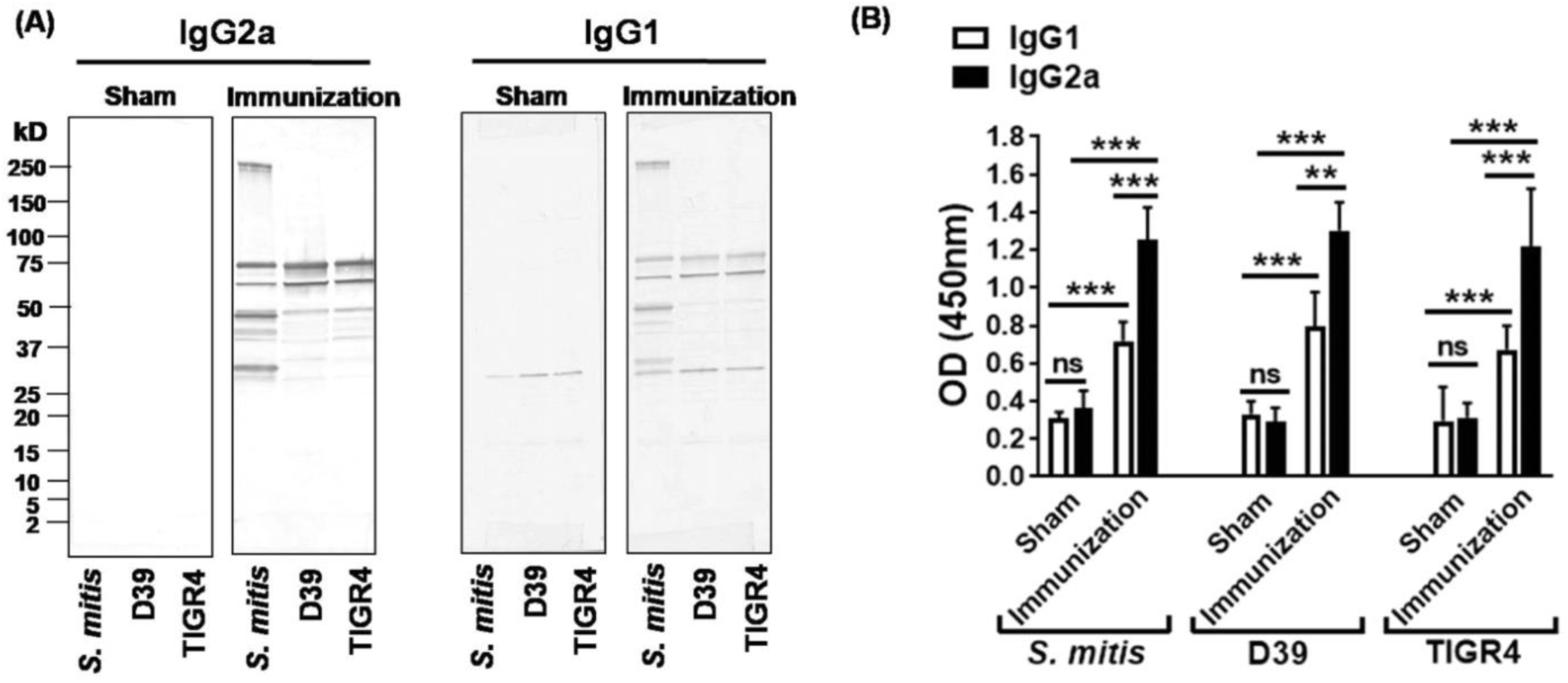
IgG2a and IgG1 antibodies from the *S. mitis*-immunized mice cross-react with *S. pneumoniae.* CD1 mice were intranasally immunized with live *S. mitis* and sera were collected to analyze antibody responses to *S. pneumoniae* strains D39 (serotype 2) and TIGR4 (serotype 4). (A) Cross-reactivity of *S. mitis*-specific IgG isotypes with *S. pneumoniae* serotypes by Western blotting. Each lane was loaded with 50 μg of protein from the indicated bacterial species. The sera were diluted 1:1000. (B) Levels of antibodies reactive to *S. mitis* and *S. pneumoniae* serotypes by whole cell ELISA. The sera were diluted 1:100. Data are shown as mean ± SD and pooled from two independent experiments with 4 mice in each group. Unpaired student’s *t* test was used to compare the levels of antibodies between sham-treated and immunized mice. **p<0.01; ***p<0.001. Abbreviations: ns = not significant; D39 – *S. pneumoniae* D39; TIGR4 – *S. pneumoniae* TIGR4.

## Discussion

In this study, we have for the first time shown that serum IgG2a and IgG1 antibody isotypes induced in response to immunization of mice with *S. mitis* cross-react with *S. pneumoniae* serotypes, and that the cross-reactivity induced by IgG2a is stronger than IgG1 antibodies. Mouse IgG2a antibody is considered as a functional equivalent of human IgG1 antibodies, which constitute the most abundant among all serum IgG isotypes.^9,10^ Immunization of human infants with pneumococcal conjugate vaccines has been reported to result in rises in IgG1 antibodies and opsonophagocytosis of antibodies to *S. pneumoniae* capsular polysaccharides.^11^ In line with these findings, our current study has shown a significant increase in *S. mitis*-specific IgG2a antibodies reactive to *S. pneumoniae* serotypes 2 and 4, which hints at the implication of this antibody isotype in *S. mitis* mediated protective immunity to *S. pneumoniae*. On the other hand, mouse IgG1 antibodies, which represent the equivalent of human IgG4 antibodies that plays a role in defense against *S. pneumoniae*, have been shown to increase in response to nasopharyngeal colonization by *S. pneumoniae*.^8,12^ This is in accordance with our results that the *S. mitis* immunization gives rise to enhanced IgG1 responses reactive to *S. pneumoniae*. But the cross-reactivity induced by IgG1 antibodies was weaker than IgG2a antibodies. Future studies are required to explore which IgG isotype plays a dominant role in conferring protection against *S. pneumoniae*. Overall, our findings may be important for understanding the *S. mitis*-induced IgG2a and IgG1 responses that cross-react with *S. pneumoniae*, which may have consequences for effective pneumococcal vaccine development.

## Acknowledgements

This work was funded by the Life Science, University of Oslo, Norway and Norwegian Research Council (Grant number - 241011).

